# Identity-by-descent refines mapping of candidate regions for preaxial polydactyly in a large Chinese pedigree

**DOI:** 10.1101/139428

**Authors:** Xingyan Yang, Quankuan Shen, Xierzhatijiang Sulaiman, Hequn Liu, Minsheng Peng, Yaping Zhang

**Author notes:** Correspondence: Dr. Minsheng Peng, Kunming Institute of Zoology, Chinese Academy of Sciences, 32 Jiaochang Donglu, Kunming 650223, China, Professor Yaping Zhang, Kunming Institute of Zoology, Chinese Academy of Sciences, 32 Jiaochang Donglu, Kunming 650223, China.

## Abstract

Preaxial polydactyly (PPD) is congenital hand malformation characterized by the duplication of digit. Herein, we scan the genome-wide SNPs for a large Chinese family with PPD-II/III. We employ the refined IBD algorithm to identify the identity-by-decent (IBD) segments and compare the frequency among the patients and normal relatives. A total of 72 markers of 0.01 percentile of the permutation are identified as the peak signals. Among of them, 57markers locate on chromosome 7q36 which is associated with PPD. Further analyses refine the mapping of candidate region in chromosome 7q36 into two 380 Kb fragments within *LMBR1* and *SHH* respectively. IBD approach is a suitable method for mapping cause gene of human disease. Target-enrichment sequencing as well as functional experiments are required to illustrate the pathogenic mechanisms for PPD in the future.

## Main text

### Background

Preaxial polydactyly (PPD; OMIM#188740) is characterized as complete or partial duplication of the thumb [1]. It is one of the most common congenital deformities [2]. The worldwide incidence of PPD is 1 in 3000 births [3]. The prevalence of polydactyly in Chinese ranks third in birth defects after heart defects, central nervous system diseases [4]. Polydactyly usually occurs in complicated hereditary patterns [5]. The mainstream treatment is resection for excess digits.

A series of efforts have been performed to investigate the genetic basis for PPD. Zguricas et al conducted linkage analysis for Dutch, British, Turkish, Cuban pedigrees and mapped the candidate region to a 1.9 cM interval between D7S550 and D7S2423 of chromosome 7q36 region [6]. Heus et al. further refined the candidate region to approximately 450 Kb including five genes: *C7orf2* (i.e. *LMBR1*), *C7orf3*(i.e. *NOM1*), *C7orf4* (i.e. *LINC00244*), *HLXB9* (i.e. *MNX1*) and *RNF32* [7] by reconstructed a detailed physical map using a combination of exon trapping, cDNA selection, and EST mapping methods. Further evidence for PPD is caused by ectopic expression of *SHH* in mice, cats and humans [8]. The zone of polarizing activity regulatory sequence (ZRS), performs as the limb-specific cis-regulator, the expression of *SHH*. ZRS locates within intron 5 of the neighboring gene *LMBR1*, which is ~1 Mb upstream from *SHH* [9]. In a number of cases, mutations of ZRS disturb the expression of *SHH* at the anterior limb bud margin and consequently caused PPD [9–16]. Deletion of ZRS showed that it is both necessary and sufficient to drive expression of *SHH* in the limb bud [17]. Duplication of ZRS is unclear how this contributes to ectopic expression [9].

The common PPD only involves in hands/feet. In extreme and rare cases, PPD occur both in hands and feet. To investigate the genetic basis, Li et al. adopted a candidate gene approach to genotype nine microsatellite markers of 7q36 chromosomal region in a Chinese family with PPD both in hands and feet. By linkage analysis and haplotype construction, they located the linkaged region spanning 1.7 Mb between D7S2465 and D7D2423 [18]. It includes the 450Kb candidate region previously identified by Henus [7]. Nevertheless, the other part of genome is not investigated yet. Herein, we genotyped genome-wide SNPs and employed the identity-by-descent (IBD) to refine the mapping of potential candidate loci for PPD in the same family.

## Methods

### Patients

This study has been approved by the internal review board of Kunming Institute of Zoology, Chinese Academy of Sciences (SMKX 2012013). The six-generation pedigree (including 21 patients and 24 normal relatives) involved in this study has been described previously in Li et al [18]. All patients show hexadactyly of hands and feet. They have been diagnosed by physical examination & X-ray and assigned as isolated PPD-II on hand and isolated PPD-III on feet according to Temtamy and McKusick’s classification [19]. PPD shows autosomal dominant inheritance in this pedigree.

### SNP array

We genotyped 900015 markers in 45 individual with HumanOmniZhongHua-8 BeadChip v1.0 (Illumina). We exported the chip data in accordance with the reference sequence GRCh37 into PLINK format via GenomeStudio (Illumina). The markers on mitochondrial DNA and sex chromosomes were disregarded. We adopted a series of quality control strategies [20] by using PLINK 1.9 [21]. Two individuals with call rate < 90% were removed. The SNPs with call rate < 90%, minor allele frequency < 1%, and deviation of Hardy– Weinberg equilibrium (P<1e-6) were excluded. After filtering, a total of 595534 autosomal SNPs for 43 individuals were utilized in subsequent analyses. The data have been deposited into Dryad (XXXXXX).

### IBD detection

We used BEAGLE 4.0 [22] to phase and impute the genotype data referring to the pedigree information and the genetic map of HapMap II [23]. We detected the IBD segment with the refined IBD in BEAGLE 4.1 [24]. The IBD segment length shorter than 1cM and the logarithm of odds (LOD) score under 3 were excluded before permutation [25]. The threshold of the genome-wide significance was set to the 0.05 percentile of the distribution of the permutation p-value.

## Results

The length distribution of detected IBD segments approximates a Pareto distribution (Fig. S1). The permutation result shows the significant segments distributing widely across genomes (Fig. 1). When considering the top 0.01% outliers of signals, we find the peak signals of 72 SNPs, of which 57 markers located at 7q36 chromosomal region (Table S1). We map the markers into the IBD fragments including *LMBR1* and *SHH* (Table 1). The minimal IBD segments within *LMBR1* and *SHH* are around 380 Kb, respectively (Table S2). The IBD segments are more frequently in patient-patient (ratio; percentage) than normal-normal (ratio; percentage) (Table 2). We make annotation for the significant SNPs (Table S1). All the SNPs haven’t been reported to be associated with PPD before.

**Fig. 1.**
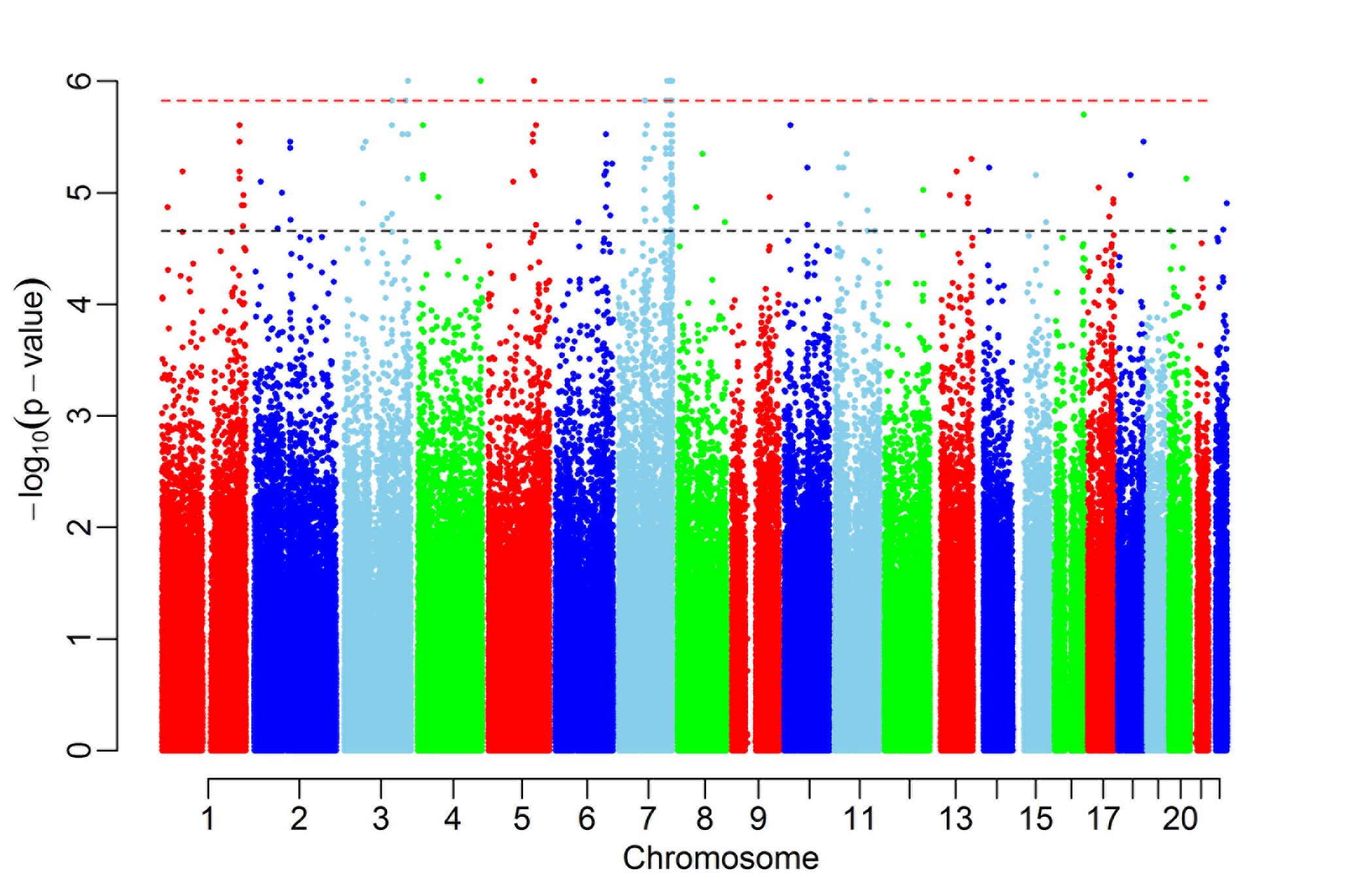
Permutation analysis after filtering out regions with low IBD sharing. The black line indicates genome-wide threshold and the red line is the 0.01 percentile of the permutation.

**Table 1.**
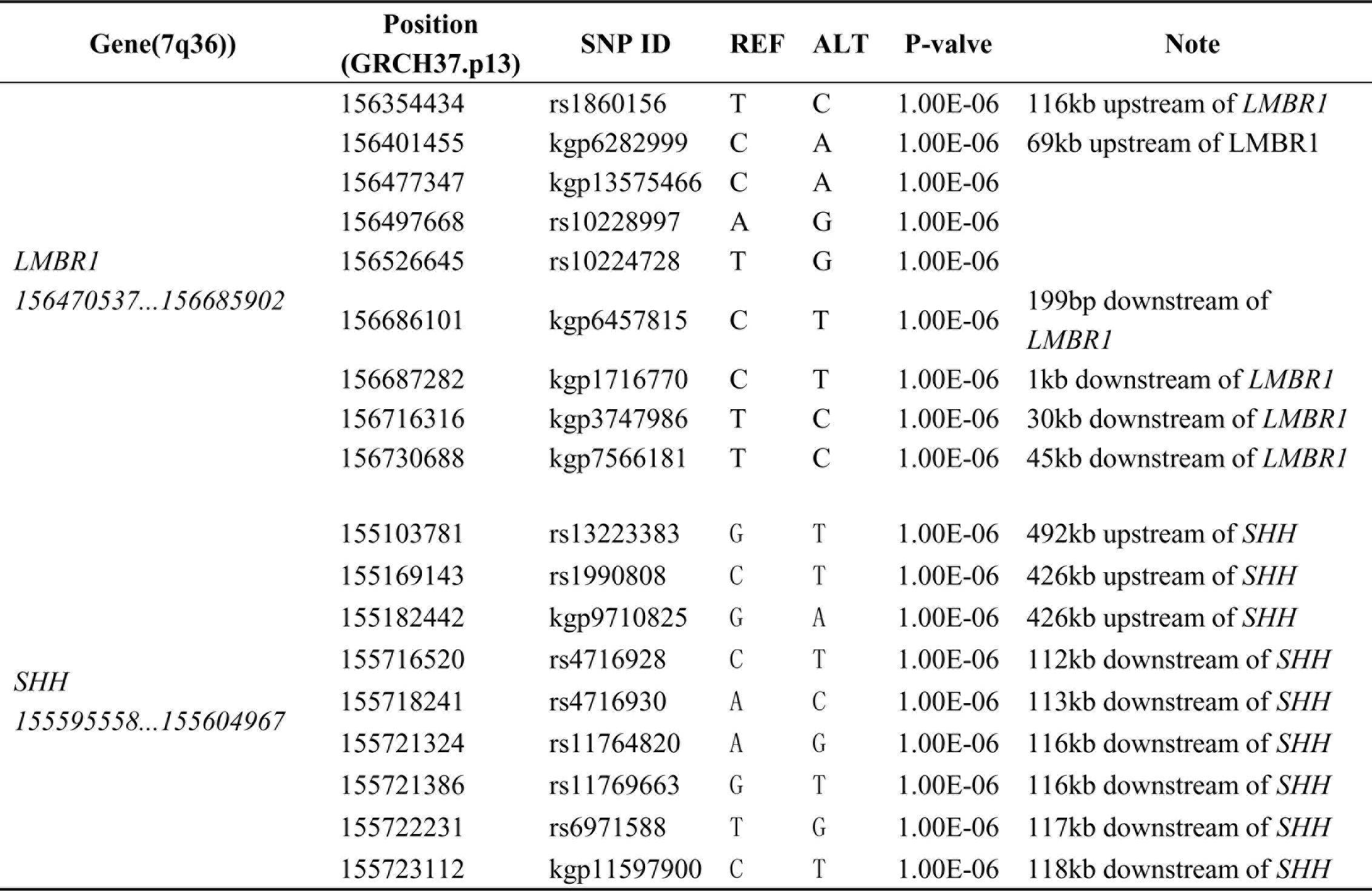
Genetic variants in the two IBD segments.

**Table 2.**
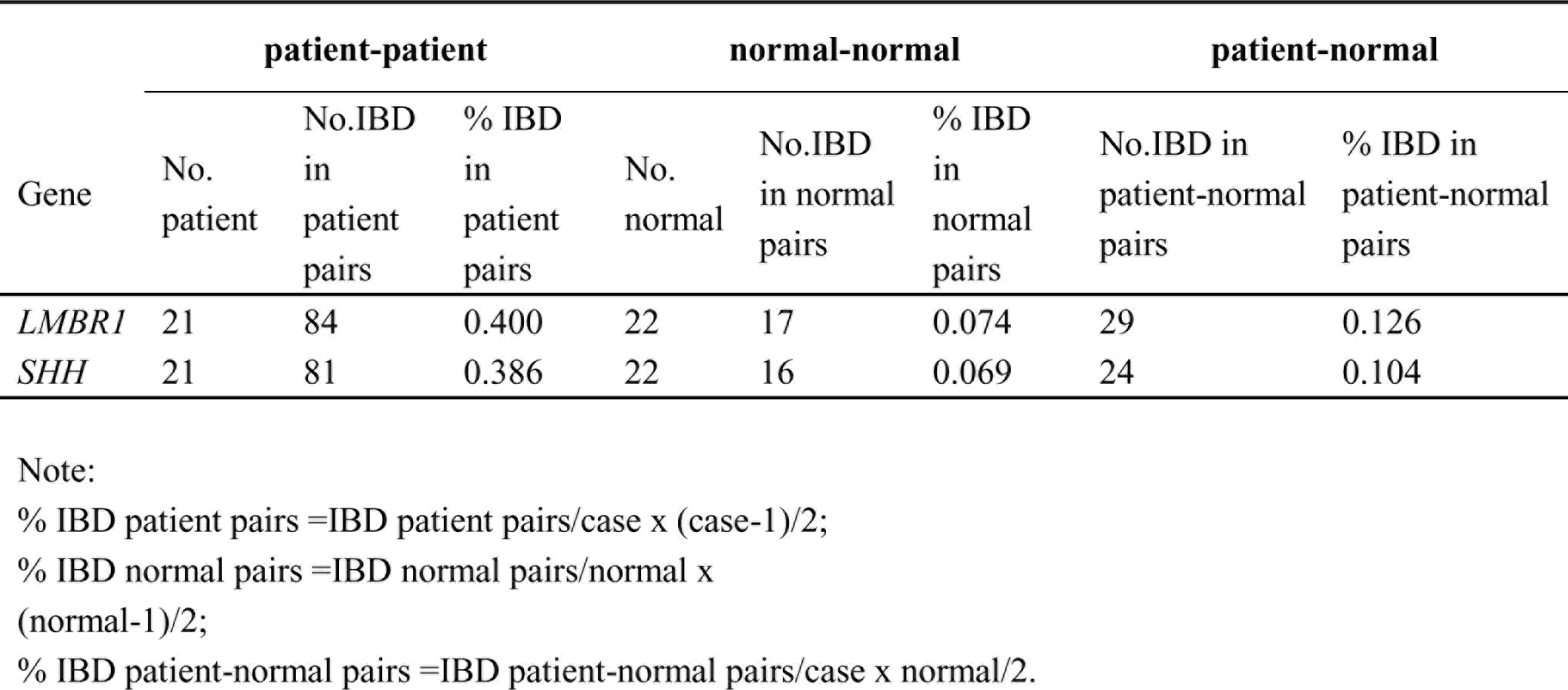
Pairwise statistics of *LMBR1* and *SHH*.

## Discussion

Our IBD analyses refine the mapping of the candidate regions for PPD into two ~380 Kb segments in 7q36 referring to *LMBR1* and *SHH* genes, respectively (Table S2). The segment for *LMBR1* includes three genes (i.e. *LMBR1*, *NOM1*, and *RNF32*) and lies within the 450Kb candidate region identified before [7]. Given that the intron 5 of *LMBR1* performed as an enhancer for *SHH* playing an important role in the pathogenesis of PPD (Table 3). Li et al. sequenced the exons of the five candidate genes and the intron 5 of *LMBR1*. But no candidate mutations were found [18]. The candidate mutations for PPD may locate in the other introns of *LMBR1*.

In addition to the segment of *LMBR1*, we also identified a segment of *SHH*. The *SHH* gene encodes sonic hedgehog, a secreted protein, which plays a key role in the limb development [26]. The ectopic expression of *SHH* in the anterior limb margin can cause PPD in mouse [27]. This has been described as well in humans [27], Hemingway cats [8] and chicken [28]. The candidate causal mutations are located in the intron 5 of *LMBR1* [9]. The duplication of ZRS can cause polysyndactyly in the Triphalangeal thumb–polysyndactyly syndrome and syndactyly type IV but not PPD [29]. Its role in PPD is unknwon. In the previous investigation of the same family, Li et al. ruled out the ZRS duplication by quantitative PCR and detected no pathogenic mutation in ZRS [18]. Consequently, the etiology of this PPD family may be another limb-specific regulatory element of *SHH* gene exists in the noncoding region. Recently, Petit et al. identified a 2 kb deletion occurring about 240 kb upstream from the *SHH* promoter in a large family with PPD-hypertrichosis. They found the 2 kb deletion repress the transcriptional activity of the *SHH* promoter in vitro [30]. Further target-enrichment sequencing and further functional experiments for *LMBR1* and *SHH* are required to identify the pathogenic mutation(s).

In summary, we refine the mapping of the candidate regions for PPD based on high-density genomic SNPs. The potential candidate mutations are most likely to locate in *LMBR1* and/or *SHH* gene. It is much improved compared with previous results [7, 18]. Our study suggests that the IBD approach is a suitable method for mapping the cause genes of human diseases. Moreover, as disruptions of topological chromatin domains can result in limb malformations [31], more attention should be paid when studying PPD in the future on this aspect.

## Abbreviations

IBD: Identity by descent
PPD: preaxial polydactyly
*LMBR1*: limb development membrane non-protein coding RNA 244
*MNX1*: motor neuron and pancreas homeobox 1
*RNF32*: ring finger protein 32

## Ethics approval and consent to participate

This study has been approved by the internal review board of Kunming Institute of Zoology, Chinese Academy of Sciences (SMKX 2012013). The patients consent to participate in this study by signing a Consent Form allowing the use of biological samples and clinical data.

## Consent for publication

A six-generation family consisting of 45 individuals including 21 affected members and 24 normal relatives was located in a rural area of Zhejiang Province, China. All patients show hexadactyly of hands and feet, diagnosed by physical examination & X-ray. According to Temtamy and McKusick’s classification it is classified as isolated PPD-II on hand and isolated PPD-III on feet. We used raw data have genotyped by Illumina HumanOmniZhongHua-8 BeadChip previously as based data for our next study. The results of the analysis of clinical data has been described previously in Li et al. Our manuscript does not contain any individual person’s data.

## Availability of data and supporting materials section

If the paper is accepted the data will be deposited into Dryad (XXXXXX).

## Authors’ contributions

XY analyzed the SNP array data and wrote the manuscript. IBD was carried out by XY and assisted by QS. XS revised the manuscript. HL performed experiments and provided patients data. MP participated in its design and revised the manuscript. All processes were guided by Dr. MP and Pro. YZ. All authors read and approved the final manuscript.

## Acknowledgments

We thank the volunteers for participating in this research. We appreciate Ping Yu, Xiaoyi Yan, Yonggang Chen, Luhang Zhao for their efforts in sampling and related information collection. We thank Nini Shi for technical assistance. This study was supported by grants from National Natural Science Foundation of China and Bureau of Science and Technology of Yunnan Province. This work was also supported by the Animal Branch of the Germplasm Bank of Wild Species, Chinese Academy of Sciences (the Large Research Infrastructure Funding). MS.P. thanks the support from the Youth Innovation Promotion Association, Chinese Academy of Sciences.

## Supplementary Figure and Table

**Fig. S1** Plot of the distribution of the IBD segments.

**Table S1** Top 0.01% peak signals.

**Table S2** IBD segments of *LMBR1* and *SHH*.

**Table S3** Mutations in intron 5 of *LMBR1*.

## References

1. Rayan G, Upton J. Congenital hand anomalies and associated syndromes. Vol. Springer Berlin Heidelberg. 2014.

2. Malik S. Polydactyly: phenotypes, genetics and classification. Clin Genet, 2014;85(3):203–12.

3. Watt J, Chung C. Duplication. Hand Clin, 2009;25(2):215–27.

4. Yan J, Huang G, Sun Y, Zhao X, Chen S, Zou S, et al. Birth defects after assisted reproductive technologies in China: analysis of 15,405 offspring in seven centers (2004 to 2008). Fertil Steril, 2011;95(1):458–460

5. Karaaslan O, Tiftikcioglu O, Aksoy M, Kocer U. Sporadic familial polydactyly. Genet Couns, 2003;14(4):401–5.

6. Zguricas J, Heus H, Morales-Peralta E, Breedveld G, Kuyt B, Mumcu F, et al. Clinical and genetic studies on 12 preaxial polydactyly families and refinement of the localisation of the gene responsible to a 1.9 cM region on chromosome 7q36. J Med Genet, 1999;36(1):32–40.

7. Heus C, Hing A, Van Baren J, Joosse M, Breedveld J, Wang C, et al. A physical and transcriptional map of the preaxial polydactyly locus on chromosome 7q36. Genomics, 1999;57(3):342–351.

8. Lettice A, Hill E, Devenney S, Hill E. Point mutations in a distant sonic hedgehog cis-regulator generate a variable regulatory output responsible for preaxial polydactyly. Hum Mol Genet, 2008;17(7):978–85.

9. Lettice A, Heaney J, Purdie A, Li L, de Beer P, Oostra A, et al. A long-range *Shh* enhancer regulates expression in the developing limb and fin and is associated with preaxial polydactyly. Hum Mol Genet, 2003;12(14):1725–1735.

10. Wieczorek D, Pawlik B, Li Y, Akarsu A, Caliebe A, May J, et al. A specific mutation in the distant sonic hedgehog (*SHH*) cis-regulator (ZRS) causes Werner mesomelic syndrome (WMS) while complete ZRS duplications underlie Haas type polysyndactyly and preaxial polydactyly (PPD) with or without triphalangeal thumb. Hum Mutat, 2010;31(1):81–9.

11. Farooq M, Troelsen T, Boyd M, Eiberg H, Hansen L, Hussain S, et al. Preaxial polydactyly/triphalangeal thumb is associated with changed transcription factor-binding affinity in a family with a novel point mutation in the long-range cis-regulatory element ZRS. Eur J Hum Genet, 2010;18(6):733–6.

12. Wang Q, Tian H, Shi Z, Zhou T, Wang Y, Shu Z, et al., A single C to T transition in intron 5 of *LMBR1* gene is associated with triphalangeal thumb-polysyndactyly syndrome in a Chinese family. Biochem Biophys Res Commun, 2007;355(2):312–7.

13. VanderMeer E, Lozano R, Sun M, Xue Y, Daentl D, Jabs W, et al. A novel ZRS mutation leads to preaxial polydactyly type 2 in a heterozygous form and Werner mesomelic syndrome in a homozygous form. Hum Mutat, 2014;35(8):945–8.

14. Semerci N, Demirkan F, Özdemir M, Biskin E, Akin B, Bagci H, et al. Homozygous feature of isolated triphalangeal thumb-preaxial polydactyly linked to 7q36: no phenotypic

15. Furniss D., et al., A variant in the sonic hedgehog regulatory sequence (ZRS) is associated with triphalangeal thumb and deregulates expression in the developing limb. Hum Mol Genet, 2008;17(16):2417–23.

16. Zhao X, Yang W, Sun M, Zhang X. ZRS mutations in two Chinese Han families featuring triphalangeal thumbs and preaxial polydactyly. Chin J Med Genet. 2016;33(3):281–5.

17. Hill E. How to make a zone of polarizing activity: insights into limb development via the abnormality preaxial polydactyly. Dev Growth Differ, 2007;49(6):439–48.

18. Li H, Wang Y, Wang X, Wu S, Yu P, Yan Y, et al. Mutation analysis of a large Chinese pedigree with congenital preaxial polydactyly. Eur J Hum Genet, 2009;17(5):604–10.

19. Temtamy A, McKusick A. The genetics of hand malformations. Birth Defects Orig Artic Ser, 1978;14(3):i–xviii, 1-619.

20. Anderson C.A., et al., Data quality control in genetic case-control association studies. Nat Protoc, 2010;5(9):1564–73.

21. Chang C, Chow C, Tellier C, Vattikuti S, Purcell M, Lee J. Second-generation PLINK: rising to the challenge of larger and richer datasets. Gigascience, 2015;4:7.

22. Browning L, Browning R. Improving the accuracy and efficiency of identity-by-descent detection in population data. Genetics, 2013;194(2):459–471.

23. Frazer A, Ballinger G, Cox R, Hinds A, Stuve L, Gibbs A, et al. A second generation human haplotype map of over 3.1 million SNPs. Nature, 2007;449(7164):851–61.

24. Browning L, Browning R. Refined IBD: a new method for detecting identity by descent in population samples. Genet Epidemiol, 2012;36(7):737–737.

25. Westerlind H, Imrell K, Ramanujam R, Myhr M, Celius G, Harbo F, et al. Identity-by-descent mapping in a scandinavian multiple sclerosis cohort. Eur J Hum Genet, 2015;23(5):688–692.

26. Kvon Z, Kamneva K, Melo S, Barozzi I, Osterwalder M, Mannion J, et al. Progressive loss of function in a limb enhancer during snake evolution. Cell, 2016;167(3):633–642.e11.

27. Hill E, Heaney J, Lettice A. Sonic hedgehog: restricted expression and limb dysmorphologies. J Anat, 2003;202(1):13–20.

28. Dunn C, Paton R, Clelland K, Sebastian S, Johnson J, McTeir L, et al. The chicken polydactyly (Po) locus causes allelic imbalance and ectopic expression of *Shh* during limb development. Dev Dyn, 2011;240(5):1163–72.

29. Sun M, Ma F, Zeng X, Liu Q, Zhao L, Wu X, et al. Triphalangeal thumb-polysyndactyly syndrome and syndactyly type IV are caused by genomic duplications involving the long range, limb-specific *SHH* enhancer. J Med Genet, 2008;45(9):589–95.

30. Petit F, Jourdain S, Holder-Espinasse M, Keren B, Andrieux J, Duterque-Coquillaud M, et al. The disruption of a novel limb cis-regulatory element of *SHH* is associated with autosomal dominant preaxial polydactyly-hypertrichosis. Eur J Hum Genet, 2016;24(1):37–43.

31. Lupiáñez G, Kraft K, Heinrich V, Krawitz P, Brancati F, Klopocki E, et al. Disruptions of topological chromatin domains cause pathogenic rewiring of gene-enhancer interactions. Cell, 2015;161(5):1012–25.

